# Parasite-mediated anorexia increases or decreases virulence evolution, depending on dietary context

**DOI:** 10.1101/401760

**Authors:** J. L. Hite, C. E. Cressler

## Abstract

Parasite-mediated anorexia is a ubiquitous, but poorly understood component of host-parasite interactions. These temporary but substantial reductions in food intake (range: 4-100%) limit exposure to parasites and alter within-host physiological processes that regulate parasite development, production, and survival, such as energy allocation, immune function, host-microbiota interactions, and gastrointestinal conditions. By altering the duration, severity, and spread of infection, anorexia could substantially alter ecological, evolutionary, and epidemiological dynamics. However, these higher-order implications are typically overlooked and remain poorly understood — even though medical (e.g., non-steroidal anti-inflammatory drugs, vaccines, targeted signaling pathways, calorie restriction) and husbandry practices (e.g., antibiotic and diet use for rapid growth, nutrient supplementation) often directly or indirectly alter host appetite and nutrient intake. Here, we develop theory that helps elucidate why reduced food intake (anorexia) can enhance or diminish disease severity and illustrates that the population-level outcomes often contrast with the individual-level outcomes: treatments that increase the intake of high quality nutrients (suppressing anorexia), can drive rapid individual-level recovery, but inadvertently increase infection prevalence and select for more virulent parasites. Such a theory-guided approach offers a tool to improve targeting host nutrition to manage disease in both human and livestock populations by revealing a means to predict how nutrient-driven feedbacks will affect both the host and parasite.

---

Parasite-mediated anorexia routinely accompanies infections in hosts ranging from insects to humans [1,2]. These substantial but temporary reductions in food intake are typically considered a negative byproduct of infection and many medical and veterinary practices (e.g., non-steroidal anti-inflammatory drugs, vaccines, antibiotic or diet use for rapid growth, and nutritional supplementation) directly or indirectly subvert anorexia [3,4]. On the other-hand, studies on calorie restriction indicate that reduced food intake during infection can improve host health and recovery [5,6]. These findings have fostered the development of novel treatment and dietary programs ranging from controlled fasting to targeting the signaling pathways that control appetite. Yet, the evolutionary and epidemiological consequences of how changing food intake (either through parasite-mediated anorexia or prescribed calorie restriction) remain overlooked and therefore poorly understood. Elucidating these key implications would provide a framework to reconcile the seemingly conflicting results from these two bodies of research and facilitate predictive ability on when and how to treat anorexia or when to recommend calorie restriction.

Doing so requires accounting for how changes in food intake affects both the host and the parasite. Parasite-mediated anorexia (and other forms of calorie restriction) may strongly impact the fitness of both hosts and parasites by, for example, reducing the ingestion of parasites [7,8], starving resident parasites of macro- and micronutrients [9,10], or limiting foods such as lipids and carbohydrates that strongly inhibit or suppress immune function [11–13]. These nutrient-dependent processes may act simultaneously with contrasting effects on host and parasite fitness, making it difficult to predict the net effect of anorexia on epidemiological dynamics or the evolution of parasites. In other words, without accounting for these complex interactions between both host and parasite, treatment plans could inadvertently select for more virulent parasites that cause more severe disease [14,15]. To date, however, most studies have focused on how anorexia alters host physiology. Surprisingly few studies have examined how anorexia affects parasite traits such as virulence (harm caused to hosts) and transmission, key traits that can evolve rapidly and routinely undermine target treatment plans (e.g., vaccines; [16–18]).

We propose a straightforward theoretical approach to this issue by integrating principles from nutritional ecology and evolutionary epidemiology. We develop a model to illustrate how theory can greatly advance our ability to predict when anorexia should benefit the host, parasite, both, or neither. We illustrate that theory is essential to: (1) estimating how various nutrient-dependent feedbacks that anorexia modulates affects the host and the parasite and (2) gaining insight into the short and long-term consequences of how changing food intake affects host health, parasite evolution, and disease dynamics. Specific nutrients are widely known to differ in the degree to which they bolster or suppress host health and immune function [9,11–13,19]. Therefore, we examined how reducing within-host nutrients, *N* (via parasite-mediated anorexia or calorie restriction) could alter parasite fitness under three dietary contexts, where nutrients can either promote or inhibit host immune function:

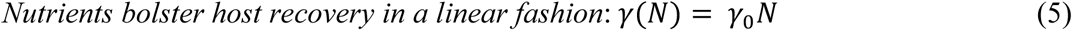

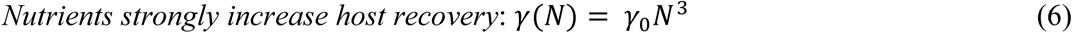

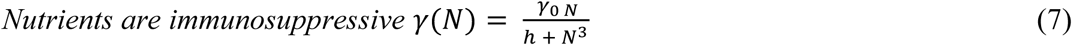

First, we consider a population of susceptible and infected hosts (*S*) and (*I*), respectively whose birth rates (*b*) and therefore population densities (*d*) change through time (*t*), depending on the level of within-host nutrients (*N*). We then introduce a free-living parasite (*Z*) into a completely susceptible host population and as a first simplifying assumption, we assume that the parasite does not evolve in response to within-host nutrients. However, because parasite transmission depends on host density, parasite-mediated anorexia which lowers within-host resources and therefore, host birth rates can still affect the expected number of secondary infections produced by a single infection i.e., parasite fitness, (*R*_0_). Therefore, we used the next-generation matrix theorem [20] to quantify how changing within-host nutrients affects parasite fitness and ability to invade or cause an epidemic, in the absence of parasite evolution. Next, we considered a case where parasites can respond to the level of nutrients within hosts (which, again anorexia modulates), by modifying production (*ϵ*) which governs the harm caused to hosts (virulence, *v*). We can use evolutionary invasion analysis [20] to access the how anorexia in infected hosts affects the evolution of parasite virulence by evaluating whether a rare mutant parasite (*Z*_*m*_) with novel production (*ϵ*_*m*_) can invade a system where *S, Q*, and *Z* remain constant.

## Results

*Parasite fitness in the absence of evolution*

We derived parasite fitness, the basic reproductive rate (*R*_0_)

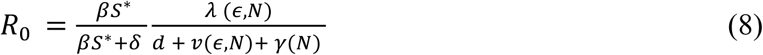

Biologically, *R*_0_ represents the ratio of parasite gains to losses or the probability that a free-living parasite infects a susceptible host 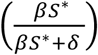, the shedding rate of parasites from infected hosts (*λ*(*ϵ,N*)), and the duration of infection 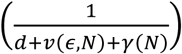, which is determined by the background death rate of hosts (*d*), the parasite-induced mortality rate (virulence, *v*(*ϵ, N*)), and host recovery rate (*γ*(*N*)). *S*\* is the equilibrium density of susceptible hosts in the absence of disease (i.e., at the disease-free boundary). Notice that within-host nutrients (*N*) (which anorexia modulates) govern three key parameters important to host fitness. Parasite fitness can increase by augmenting shedding or prolonging infection via reduced host recovery or host mortality.

The model indicates that, in the absence of evolution, regardless of dietary context, nutrient-dependent feedbacks create a unimodal (hump-shaped) relationship between food intake and parasite fitness, (*R*_0_). The implication of this pattern is that suppressing anorexia by increasing feeding will increase parasite fitness (if anorexia is extreme) or decrease it (if anorexia is mild). Taking the parasite’s perspective, if hosts typically have high ingestion, then anorexia will always increase parasite fitness. However, the different effects of nutrients on immunity carry important consequences for which parasite strains have the highest fitness. With diets that either weakly or strongly promote host recovery (Eq. 5, Fig. 2a, and Eq. 6; Fig. 2b, respectively), low virulence strains always have higher fitness than more virulent strains; moreover, suppressing anorexia can even drive some strains extinct (*R*_0_ < 1), particularly the most virulent strains, which exhibit sharper declines at high nutrient levels. Conversely, with immunosuppressive diets, high virulence strains will have higher fitness than low virulence strains when nutrient levels are low (Fig. 2c). Together, these results underscore that how anorexia affects parasite fitness and virulence depends sensitively on dietary context.

**Fig. 1.**
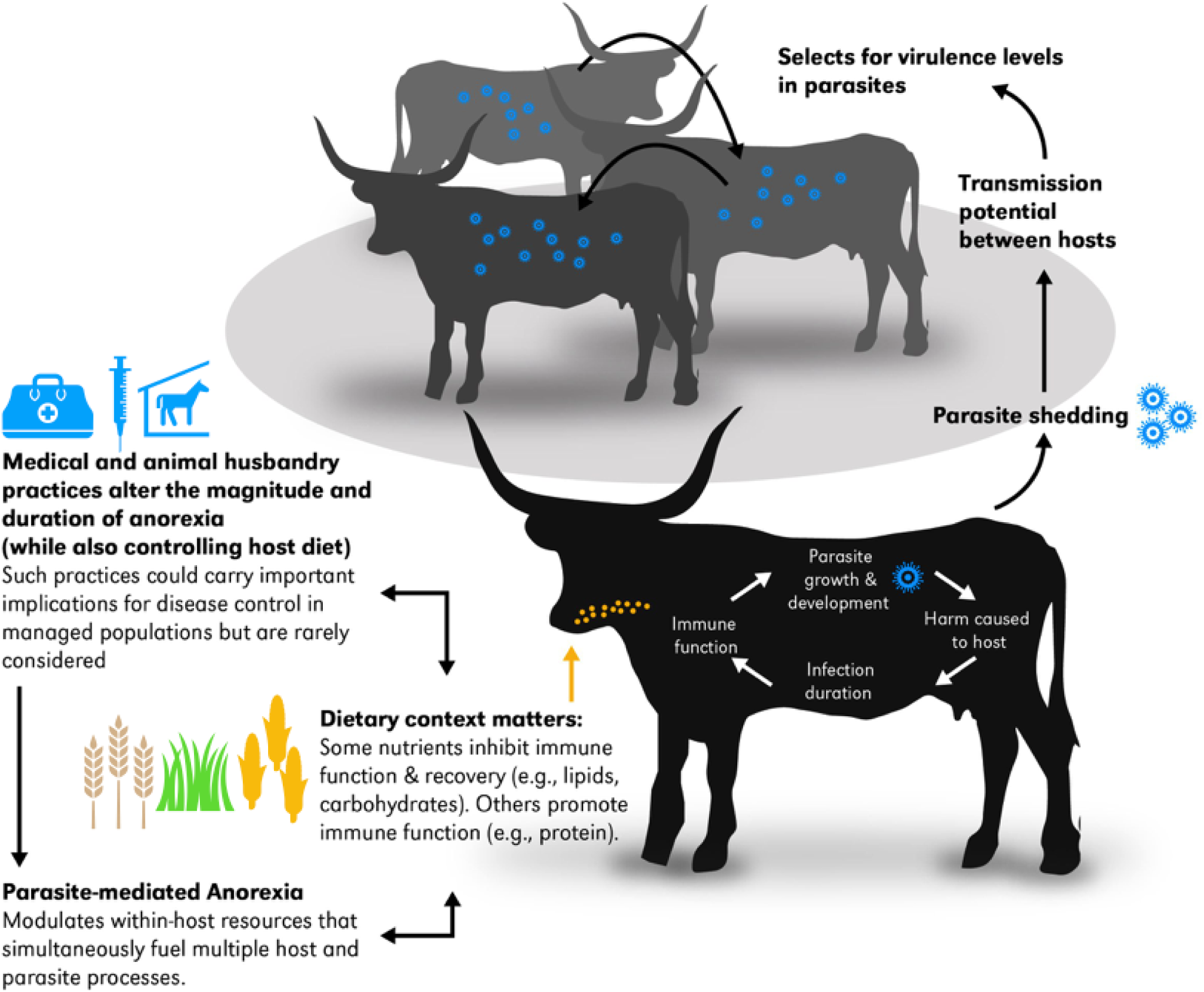
Schematic illustrating how changes in parasite-mediated anorexia alters within-host nutrients, host immune function, and parasite production under different dietary contexts. Anorexia and dietary context modulate complex interactions at the individual-level could scale-up to drive epidemiological and evolutionary outcomes with implications for managing human, livestock, and wildlife diseases.

**Fig. 2.**
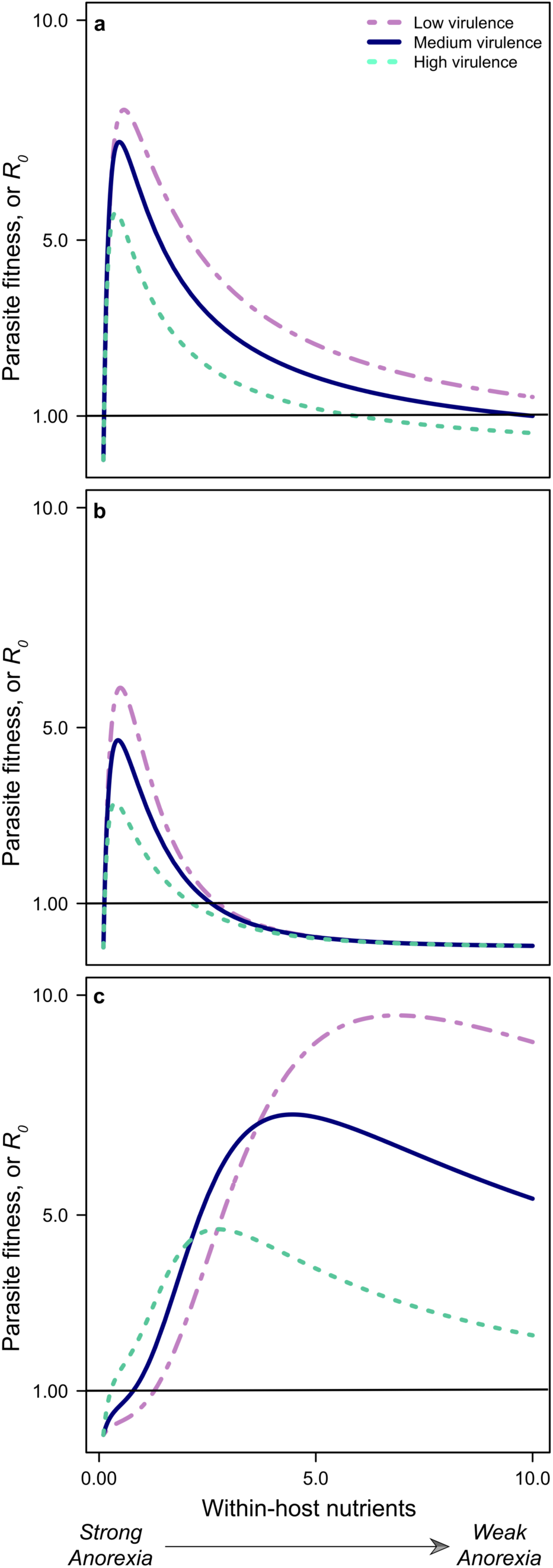
Changes in parasite-mediated anorexia alter parasite fitness (R_0_) in the absence of evolution. The figure illustrates how food intake and dietary context effects parasite strains that differ in virulence levels: Calorie intake increases host immune function and promotes recovery with either (*a*) ‘weakly’ (*γ* = *γ*_*0*_N) or (*b*) ‘strongly’ (*γ* = *γ*_*0*_*N*^3^) or (*c*) calorie intake interferes with host recovery 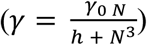. Parasites can invade above the solid horizontal line (i.e., *R*0 > 1).

### The evolution of parasite virulence

Next, we examined how parasite-mediated anorexia could alter the evolution of virulence by considering cases where within-host resource conditions strongly affect parasite evolution and disease dynamics as shown, for example, in malaria [21] and bacteria in zooplankton and managed fish populations [22,23]. Again, we derived the invasion fitness of the mutant parasite (*r*_*m*_) using the next generation theorem, which is the expected lifetime transmission from a host infected with a rare mutant:

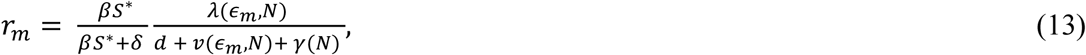

where *S*\* is now the number of hosts left uninfected by the resident parasite. Any mutant parasite with *r*_*m*_ > 1 will increase from rarity and is assumed to outcompete and replace the resident parasite. The goal is to find evolutionarily stable (ES) values of the evolving trait, *ϵ*_*m*_ = *ϵ*\*. At such ES values, the selection gradient (the derivative of *r*_*m*_ with respect to the evolving trait *ϵ*_*m*_) vanishes and invasion fitness is maximized. Any root of the selection gradient will be evolutionarily stable (ES) if the costs of parasite production increase faster than the benefits (e.g.,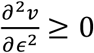 and 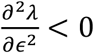 see S1 Appendix). Such roots will satisfy:

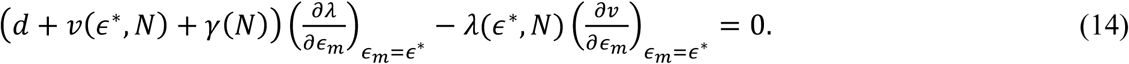

Using the biological intuition behind the terms of the invasion fitness expression, the ES production (*ϵ*\*) will satisfy:

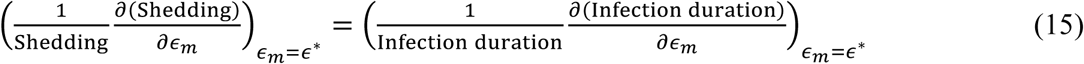

That is, at the ES production, the relative increase in shedding from increasing production is perfectly balanced by an equal decrease in infection duration. This balance depends on tension between nutrients fueling host recovery (which prolongs host life span and thus the duration of infection) and nutrients fueling parasite shedding. These nutrient-driven dynamics exert contrasting forces on parasite fitness and therefore, the evolution of virulence.

To explore how increasing within-host nutrients affected the ES level of parasite production, *ϵ*\*, we evaluated how anorexia affects parasite evolution by implicitly differentiating the fitness gradient expression (Eq. 14) with respect to *N*. In brief, the crucial points of this complex expression highlight the tension between the effects of nutrients on recovery 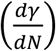, shedding 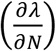,infection duration (*d* + *v*(*ϵ*\*, *N*) + *γ*(*N*)), and virulence 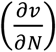; See S1 Appendix for expanded explanations. Existing theory predicts that any factor that shortens infection duration (e.g., increased recovery) should promote the evolution of increased virulence [24–26]. Yet, our results indicate that even if nutrients increase recovery rate 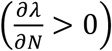, ES production (and hence, virulence) may still decrease, depending on how sensitively parasite shedding depends on within-host nutrients. For example, given our functions for virulence and shedding, with nutrients that moderately increase host recovery (*γ* = *γ*_0_ N, increasing host food intake (suppressing anorexia), can drive a decrease in ES production with a subsequent increase in virulence (Fig. 3 small-dotted black lines). Whereas if nutrients strongly promote host recovery (*γ* = *γ*_0_*N*^3^), increasing food intake drives a marked increase in both ES production and virulence (Fig. 3 solid cyan lines). Counterintuitively, when nutrients inhibit host recovery 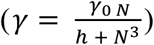, increasing food intake decreases both ES production and virulence (Fig. 3 large-dashed magenta lines). Hence, with nutrients that inhibit immune function, host that exhibit strong anorexia could select for higher parasite production and increased virulence, driving more severe disease dynamics.

**Fig. 3.**
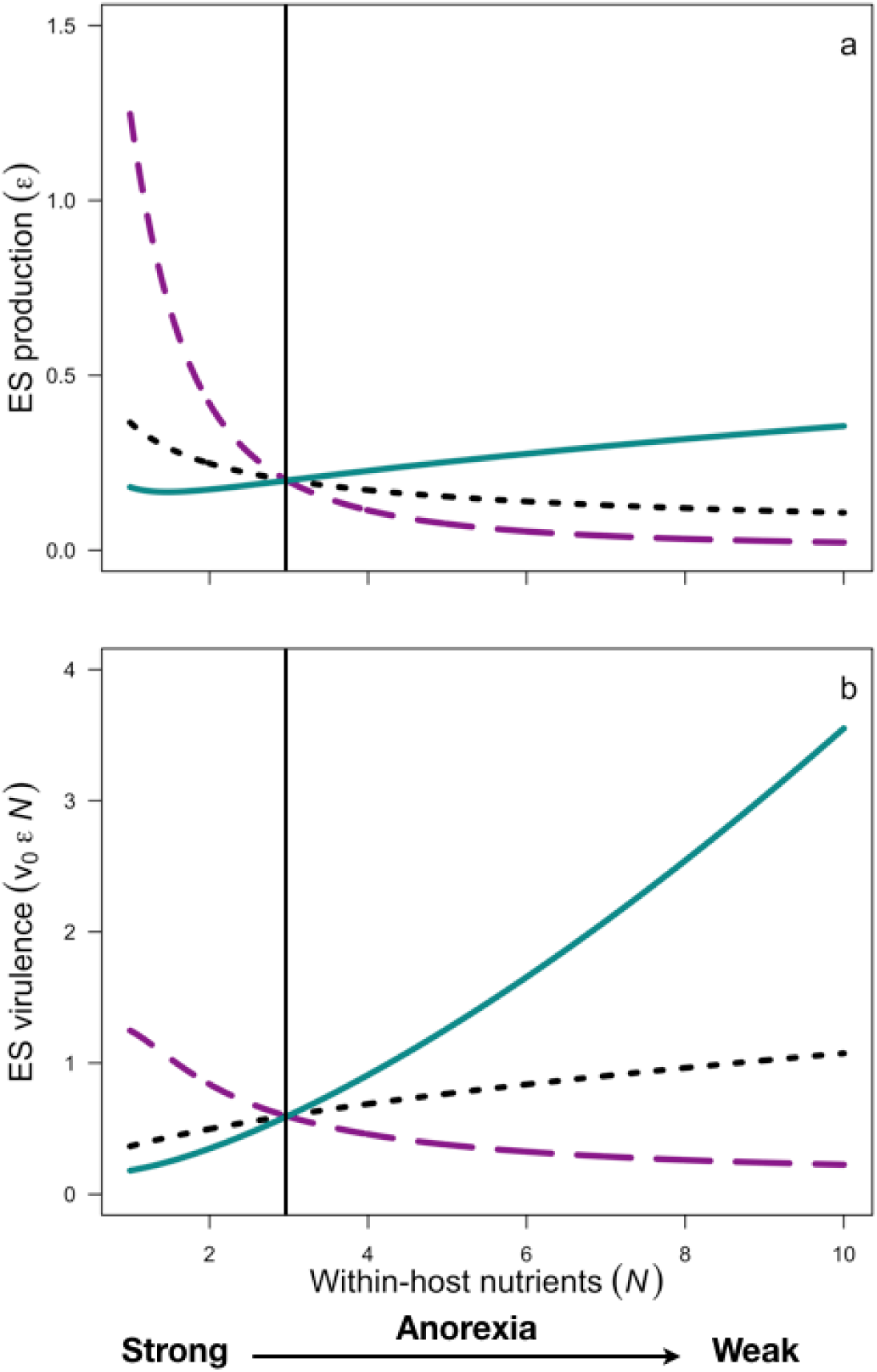
Evolutionary consequences of changing the magnitude of parasite-mediated anorexia under different dietary contexts. With nutrients that moderately increase host recovery (*γ* = *γ*_*0*_N) increasing host food intake (suppressing anorexia), can drive a decrease in ES production with a subsequent increase in virulence (Fig. 3 small-dotted black lines). Whereas if nutrients strongly promote host recovery (*γ* = *γ*_*0*_*N*^3^), increasing food intake drives a marked increase in both ES production and virulence (Fig. 3 solid cyan lines). Counterintuitively, when nutrients inhibit host recovery 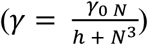, increasing food intake leads to a decrease in both ES production and virulence (Fig. 3 large-dashed magenta lines). Hence, with nutrients that inhibit immune function, strong host anorexia can select for higher parasite production and increased virulence, driving more severe disease dynamics. Solid vertical line represents a host’s ‘baseline’ level of food intake, imagining scenarios where medical interventions shift this baseline.

In addition to altering parasite evolution, changes to food intake (within-host nutrients) altered disease prevalence and harm to host populations. When nutrients increase host recovery, increasing food intake (suppressing anorexia) decreases infection prevalence, regardless of whether the parasite evolves in response to within-host nutrients (Fig. 4a-k). However, with immunosuppressive nutrients, increasing food intake can either decrease or increase infection prevalence, depending critically on how strongly within-host nutrients affects parasite fitness and evolution (Fig. 4 dotted vs. solid lines). Notably, these population-level effects (Fig. 4) often contrast with the individual-level effects (Fig. 3). Consequently, treatments that increase food intake (e.g., by increasing host appetite) with high quality nutrients could drive rapid individual-level recovery, but inadvertently increase infection prevalence and disease severity at the population-level.

**Fig. 4.**
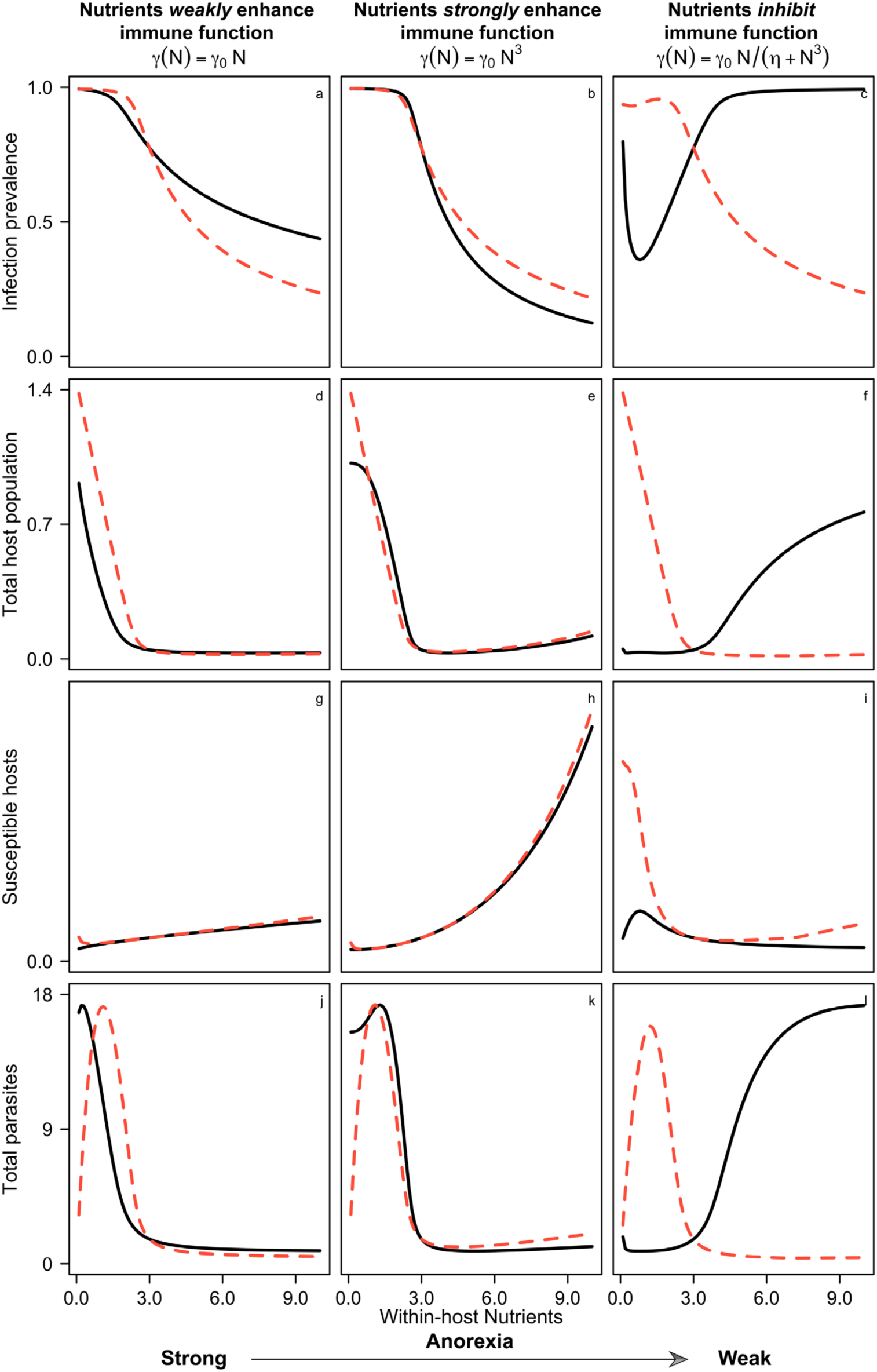
Epidemiological and population-level consequences of changing within-host nutrients (e.g., via parasite-mediated anorexia or calorie restriction) under different dietary contexts with (solid lines) and without (dashed lines) parasite evolution. With diets that increase host recovery, suppressing anorexia (within-host nutrients increase), always decreases infection prevalence, regardless of whether parasite evolution occurs. Conversely, when nutrient intake inhibits immune function and host recovery, predictions for infection prevalence with or without parasite evolution vary substantially; with parasite evolution, strong anorexia reduces infection prevalence and suppressing anorexia can increase infection prevalence. This bimodal relationship emerges from tension between two forces that pull in contrasting directions; nutrients suppress immune function and host recovery but fuel parasite development and growth.

## Discussion

### Potential adaptive significance of anorexia and implications for the disease management

The ubiquity of parasite-mediated anorexia, combined with increased appreciation for the short-term benefits of managed calorie restriction on host health, and the wide-spread use of many biomedical practices that alter food intake, have led to an increasing need to identify contexts where anorexia benefits the host, parasite, both, or neither [1,27,28]. A key merit of a theory-guided approach is that it reveals the contrasting effects of within-host nutrients on parasites and hosts. The outcomes of parasite-mediated anorexia on host health and disease severity not only depend on dietary context but are also likely modulated by nutrient-driven tension between hosts *and* parasites (even though most studies on calorie restriction focus solely on the host).

This method also yields a straightforward method for integrating empirically-derived parameters to quantify the effects of anorexia on parasite fitness and disease severity (at both individual and population levels), thus identifying key points for intervention. For example, if parasite production depends sensitively on within-host nutrients, and host diet promotes host recovery but strongly suppresses parasite production, medical- or management-driven changes to ‘background’ levels of food intake could drive the evolution of more virulent parasites (Figs. 2,3) and lead to counterintuitive increases in infection prevalence (Fig. 4a-c). At first glance, this result is unsurprising; any factor that shortens infection duration or reduces parasite-driven mortality (e.g., increased recovery due to vaccination) should promote the evolution of increased virulence [24–26]. Indeed, such findings have fostered the development of more integrated approaches to vaccine and antibiotic development to reduce unintended threats to public health. However, while dietary protocols represent integral components to many medical (e.g., vaccines, calorie restriction, intravenous nutrient administration) [29,30], and farming practices (rapid biomass production via high-calorie or high-fat diets, low-level antibiotics to promote growth, nutrient supplementation), the epidemiological or evolutionary consequences of these practices are rarely quantified or considered [9]. Finally, our results show that this approach can reveal previously overlooked and unintended aspects of anorexia on epidemiological and evolutionary consequences, producing new hypotheses for disease control with potential applications toward human, livestock, and wildlife health [15,31,32].

Our results also carry implications for disease control in managed populations. Current animal husbandry practices intensively select for growth (through both genetic selection and diet) and these lines often experience more severe infection characteristics and loss of appetite relative to animals from slower-growing lines [33,34]. Our results suggest that emphasizing rapid biomass production and growth, combined with host diet and treatments that directly or indirectly alter host calorie intake, could backfire; overfeeding hosts could improve individual-level recovery but promote more virulent parasites, subsequently driving the evolution of higher infection prevalence and more severe disease at the population level. Incorporating these interactions into current management practices could potentially advance current efforts to reduce demand for certain medical interventions (e.g., anthelminthics, antibiotics), and help slow the evolution of resistance.

### Materials and Methods

Our model for host-parasite dynamics is as follows:

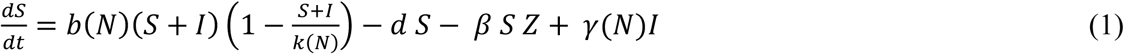

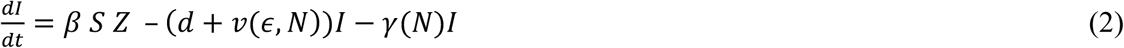

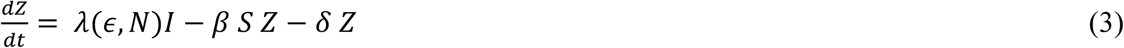

All hosts are born susceptible with per-capita birth rate *b*(*N*) and the strength of density-dependence *k*(*N*) dependent on within-host nutrient levels, *N*. Background mortality rate (*d*) is independent of density and nutrients. Hosts become infected through contact with free-living parasites (*Z*) at rate, *β Z*. Infected hosts die due to infection (virulence) at rate *v*(*ϵ, N*) = *v*_*0*_*ϵN*, with *v*_*0*_ the per-parasite virulence and shed free-living parasites into the environment at rate, *λ*(*ϵ, N*). Shedding links to parasite production via a saturating relationship[31],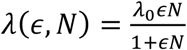, with *λ*_*0*_ the maximum shedding rate. Hence, both virulence and shedding rates depend on parasite production (*ϵ*), which depend linearly on within-host nutrients (*N*). Free-living pathogens die at a background rate, *δ*. Infected hosts recover at rate, *γ*(*N*), which, again, may increase or decrease with nutrients. Model analysis was carried out using MATHEMATICA 11.1[35] (S1 Appendix).

The expanded model, including hosts infected with the mutant parasite (*I*_*m*_) is:

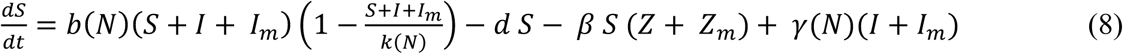

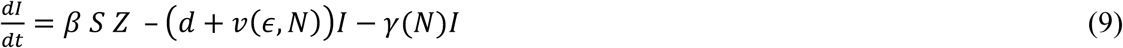

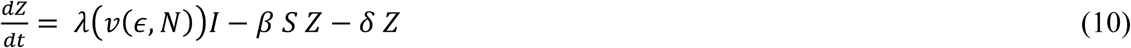

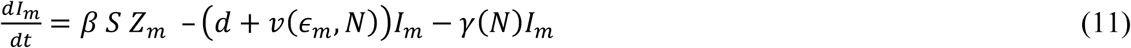

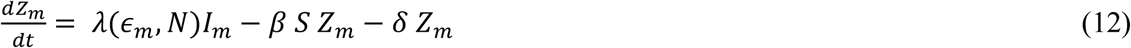

## Acknowledgements

We thank Kristi Montooth and Justin Buchanan for insightful discussions.

## Author contributions

JLH and CEC conceived and implemented the study. JLH wrote the first draft of the manuscript. Both JLH and CEC

## Supplementary Materials

Materials and Methods

Mathematica Code

## References and Notes

1. Hart BL. Behavioral adaptations to pathogens and parasites: Five strategies. Neurosci Biobehav Rev. 1990;14. doi:10.1016/S0149-7634(05)80038-7.

2. Ayres JS, Schneider DS. The Role of Anorexia in Resistance and Tolerance to Infections in Drosophila. PLOS Biol. Public Library of Science; 2009;7: e1000150. Available: https://doi.org/10.1371/journal.pbio.1000150

3. Fox MT, Uche UE, Vaillant C, Ganabadi S, Calam J. Effects of Ostertagia ostertagi and omeprazole treatment on feed intake and gastrin-related responses in the calf. 2002;105: 285–301.

4. Sharon KP, Duff GC, Paterson JA, Dailey JW, Carroll JA, Marceau EA. Case Study: Effects of timing of a modified-live respiratory viral vaccination on performance, feed intake, antibody titer response, and febrile response of beef heifers. Prof Anim Sci. 2013;29: 307–312. doi: https://doi.org/10.15232/S1080-7446(15)30237-0.

5. Wang A, Huen SC, Luan HH, Yu S, Zhang C, Gallezot J-D, et al. Opposing Effects of Fasting Metabolism on Tissue Tolerance in Bacterial and Viral Inflammation. Cell. United States; 2016;166: 1512–1525.e12. doi:10.1016/j.cell.2016.07.026.

6. Cheng C-W, Villani V, Buono R, Wei M, Kumar S, Yilmaz OH, et al. Fasting-Mimicking Diet Promotes Ngn3-Driven beta-Cell Regeneration to Reverse Diabetes. Cell. United States; 2017;168: 775–788.e12. doi:10.1016/j.cell.2017.01.040.

7. Hite JL, Penczykowski RM, Shocket MS, Griebel K, Strauss AT, Duffy MA, et al. Allocation, not male resistance, increases male frequency during epidemics: A case study in facultatively sexual hosts. Ecology. 2017; doi:10.1002/ecy.1976.

8. Grenfell BT. Gastrointestinal Nematode Parasites and the Stability and Productivity of Intensive Ruminant Grazing Systems. Philos Trans R Soc B Biol Sci. 1988;321: 541–563. doi:10.1098/rstb.1988.0107.

9. Kyriazakis I, Tolkamp BJ, Hutchings MR. Towards a functional explanation for the occurrence of anorexia during parasitic infections. Anim Behav. 1998;56. doi:10.1006/anbe.1998.0761.

10. Povey S, Cotter SC, Simpson SJ, Wilson K. Dynamics of macronutrient self-medication and illness-induced anorexia in virally infected insects. J Anim Ecol. 2014;83. doi:10.1111/1365-2656.12127.

11. Povey S, Cotter SC, Simpson SJ, Wilson K. Dynamics of macronutrient self-medication and illness-induced anorexia in virally infected insects. Altizer S, editor. J Anim Ecol. 2014;83: 245–255. doi:10.1111/1365-2656.12127.

12. Singer MS, Mason PA, Smilanich AM. Ecological immunology mediated by diet in herbivorous insects. Integr Comp Biol. 2014;54. doi:10.1093/icb/icu089.

13. Mason AP, Smilanich AM, Singer MS. Reduced consumption of protein-rich foods follows immune challenge in a polyphagous caterpillar. J Exp Biol. 2014;217: 2250 LP- 2260. Available: http://jeb.biologists.org/content/217/13/2250.abstract

14. Cressler CE, Nelson WA, Day T, Mccauley E. Disentangling the interaction among host resources, the immune system and pathogens. Ecol Lett. 2014;17: 284–293. doi:10.1111/ele.12229.

15. Hite JL, Cressler CE. Resource-driven changes to host population stability alter the evolution of virulence and transmission. Philos Trans R Soc London B. 2018;

16. Gandon S, Day T. Evidences of parasite evolution after vaccination. Vaccine. 2008;26: C4–C7. doi: https://doi.org/10.1016/j.vaccine.2008.02.007.

17. Barclay VC, Sim D, Chan BHK, Nell LA, Rabaa MA, Bell AS, et al. The evolutionary consequences of blood-stage vaccination on the rodent malaria Plasmodium chabaudi. PLoS Biol. 2012;10: e1001368. doi:10.1371/journal.pbio.1001368.

18. Read AF, Baigent SJ, Powers C, Kgosana LB, Blackwell L, Smith LP, et al. Imperfect vaccination can enhance the transmission of highly virulent pathogens. PLoS Biol. 2015;13: 1–18. doi:10.1371/journal.pbio.1002198.

19. Adamo SA, Bartlett A, Le J, Spencer N, Sullivan K. Illness-induced anorexia may reduce trade-offs between digestion and immune function. Anim Behav. Elsevier Ltd; 2010;79: 3–10. doi:10.1016/j.anbehav.2009.10.012.

20. Hurford A, Cownden D, Day T. Next-generation tools for evolutionary invasion analyses. J R Soc Interface. 2010;7: 561–571. doi:10.1098/rsif.2009.0448.

21. Wale N, Sim DG, Jones MJ, Salathe R, Day T, Read AF. Resource limitation prevents the emergence of drug resistance by intensifying within-host competition. Proc Natl Acad Sci. 2017; Available: http://www.pnas.org/content/early/2017/12/11/1715874115.abstract

22. Vale PF, Salvaudon L, Kaltz O, Fellous S. The role of the environment in the evolutionary ecology of host parasite interactions. Infect Genet Evol. 2008;8: 302–305.

23. Kinnula H, Mappes J, Valkonen JK, Pulkkinen K, Sundberg L-R. Higher resource level promotes virulence in an environmentally transmitted bacterial fish pathogen. Evol Appl. Wiley/Blackwell (10.1111); 2017;10: 462–470. doi:10.1111/eva.1246.

24. Choo K, Williams PD, Day T. Host mortality, predation, and the evolution of parasite virulence. Ecol Lett. 2003; 310–315.

25. Gandon S, Mackinnon MJ, Nee S, Read AF. Imperfect vaccines and the evolution of pathogen virulence. Nature. 2001;414: 751–6. doi:10.1038/414751.

26. van Baalen M. Coevolution of recovery ability and virulence. Proceedings Biol Sci. England; 1998;265: 317–325. doi:10.1098/rspb.1998.029.

27. Sylvia KE, Demas GE. A Return to Wisdom: Using Sickness Behaviors to Integrate Ecological and Translational Research. Integr Comp Biol. England; 2017;57: 1204–1213. doi:10.1093/icb/icx05.

28. Rao S, Schieber AMP, O’Connor CP, Leblanc M, Michel D, Ayres JS. Pathogen-mediated inhibition of anorexia promotes host survival and transmission. Cell. Elsevier; 2017;168: 503–516.e12. doi:10.1016/j.cell.2017.01.00.

29. Lucht JM, Mauch-Mani B, Steiner HY, Metraux JP, Ryals J, Hohn B. Pathogen stress increases somatic recombination frequency in Arabidopsis. Nat Genet. 2002;30: 311–314. doi:10.1038/ng84.

30. Schetz M, Casaer MP, Van den Berghe G. Does artificial nutrition improve outcome of critical illness? Crit Care. England; 2013;17: 302. doi:10.1186/cc1182.

31. Hall SR, Simonis JL, Nisbet RM, Tessier AJ, Cáceres CE. Resource ecology of virulence in a planktonic host-parasite system: an explanation using dynamic energy budgets. Am Nat. 2009;174: 149–62. doi:10.1086/60008.

32. Becker DJ, Streicker DG, Altizer S. Linking anthropogenic resources to wildlife–pathogen dynamics: a review and meta-analysis. Ecol Lett. 2015;18: 483–495. doi:10.1111/ele.1242.

33. Kyriazakis I, Doeschl-Wilson A. Anorexia during infection in mammals: variation and its sources. Voluntary feed intake in pigs. Wageningen: Wageningen Academic Publishers; 2009. pp. 307–321.

34. Zaralis K, Tolkamp BJ, Houdijk JGM, Wylie ARG, Kyriazakis I. Consequences of protein supplementation for anorexia, expression of immunity and plasma leptin concentrations in parasitized ewes of two breeds. Br J Nutr. England; 2009;101: 499–509. doi:10.1017/S000711450802401.

35. Mathematica 11.1. Wolfram Res Inc. 2017;

